# YGEP: An Integrated Web Platform for Genomic Evaluation and Intelligent Breeding in Yak

**DOI:** 10.64898/2026.09.29.755494

**Authors:** Yuanyuan Yu, Zemin Li, Xinru Shan, Yangyang Fu, Yujiao Fu, Zehao Zhang, Xingyu Guo, Shikai Wang, Jiahong Zhao, Jincheng Zhong, Langda Suo, Daoliang Lan, Jiabo Wang

## Abstract

**Background:** The yak (*Bos grunniens*) is an essential livestock species in high-altitude alpine ecosystems, serving key roles in meat, milk, and wool production as well as ecological maintenance. However, traditional yak breeding systems face several limitations, including fragmented herd data management, inefficient phenotypic collection, missing pedigree records, and high technical barriers to applying genomic technologies. These factors constrain the widespread application of modern genetic evaluation methods in yak breeding programs.

**Description:** We developed the Yak Genomic Evaluation Platform (YGEP, https://yakbreeding.swun.edu.cn/en/), an integrated digital system for yak genetic selection and herd management. YGEP integrates three core analytical modules: pedigree reconstruction, genotype imputation and genomic breeding value estimation. Benchmarking on 706 yaks showed that GBLUP achieved a prediction accuracy of r = 0.96 for cannon bone circumference (CCB), and KBeagle reached 84.0% imputation accuracy with the combined reference panel.

**Conclusion:** YGEP bridges the gap between advanced genomic analytics and practical yak breeding. By integrating pedigree reconstruction, genotype imputation, and GEBV estimation into a user-friendly cloud platform, YGEP effectively eliminates technical and data bottlenecks. Overall, it provides an accessible, scalable, and standardized solution to advance digital herd management and precision genetic improvement in alpine livestock.

## 1. Background

The yak (*Bos grunniens*) is an important indigenous domestic animal that is adapted to the high-altitude ecological environment. It is also a crucial resource for meat, milk, and wool production, as well as for maintaining the ecosystem in the Qinghai-Xizang Plateau region [1,2]. Through long-term adaptation to extreme environments such as low oxygen, low temperature, and resource limitations, the yak has developed a unique genetic structure and environmental adaptation mechanism [3,4]. Therefore, analyzing the genetic resource characteristics of the yak and establishing an efficient breeding system are of great significance for promoting the sustainable development of plateau animal husbandry. With the increasing demand for efficient and precise genetic improvement, there is a growing need for breeding strategies that can make better use of the limited genetic and phenotypic resources available in yak populations.

Genetic improvement in yaks is currently constrained by limited breeding data and genomic resources. Conventional breeding relies heavily on complete pedigree and long-term phenotypic records, which are often incomplete in yak populations. These limitations reduce the efficiency of traditional selection and hinder the implementation of systematic breeding programs.

Genomic selection (GS) provides an effective alternative by using genome-wide SNP information to predict genomic estimated breeding values (GEBVs), and its feasibility has been demonstrated for growth and other important traits in yak [5,6]. However, the practical application of genomic breeding remains challenged by unreliable pedigree information and the high cost of high-density genotyping. Genotype imputation using low-density marker panels provides a cost-effective approach to increase marker density and support genomic prediction[7,8]. Together, GS and genotype imputation provide promising strategies for improving the efficiency and scalability of yak breeding. However, their practical implementation requires analytical frameworks capable of integrating genotype, phenotype, pedigree, and population information rather than relying on isolated tools.

Existing genomic resources alone are not sufficient to support the practical implementation of genomic breeding in yaks. In recent years, with the development of high-throughput sequencing technology, multi-omics analysis methods, and bioinformatics technology, several yak genetic resource databases have been established successively, such as the Yak Genome Database (YGD) and the multi-omics database[9,10]. These databases provide important data resources for the analysis of the genomic structure of the yak, the mining of functional genes, and the study of high-altitude adaptation mechanisms. For example, YGD provides access to genome sequences, gene annotations, genetic variants, and comparative genomic information, while recent multi-omics resources have further incorporated transcriptomic, proteomic, and epigenomic data to facilitate systematic investigations of yak biology and adaptation. However, the existing databases related to yaks mainly focus on the storage, retrieval, and sharing of genomic data, and their functions are usually limited to static data resource services, lacking an interactive analysis environment that supports user-defined data uploads, online computational analysis, and breeding decision assistance. Therefore, these platforms do not directly meet the comprehensive needs of modern yak breeding practices for data management, genetic assessment, and breeding decision-making [9,11].

This gap between genomic resource databases and user-oriented genomic evaluation represents an important barrier to the practical application of genomic breeding in yak. Recent tools such as IP4GS and HEGS have significantly simplified genomic selection in crop science and multi-institution collaboration [12,13]. However, IP4GS focuses on crop breeding workflows, and HEGS addresses secure multi-institution collaboration; neither targets the specific challenges of yak genomic evaluation, such as incomplete pedigrees and genetically differentiated populations. Building on these advances, yak genomic evaluation presents distinct operational requirements, such as handling incomplete pedigrees, varying marker densities, and multi-source phenotypic data. Therefore, extending integrated platform concepts to yak-specific scenarios is essential for practical genomic breeding.

To address these issues, this study developed the Yak Genomic Evaluation Platform (Yak Genomic Evaluation Platform, YGEP). This platform integrates functional modules such as population information management and genomic analysis, providing users with an integrated service process from data upload, automatic analysis to result output **(Fig. 1)**. It provides data support for cross-group genetic assessment and breeding analysis, and offers technical support for the protection of yak genetic resources, precise breeding, and the development of modern agriculture.

**Fig. 1.**
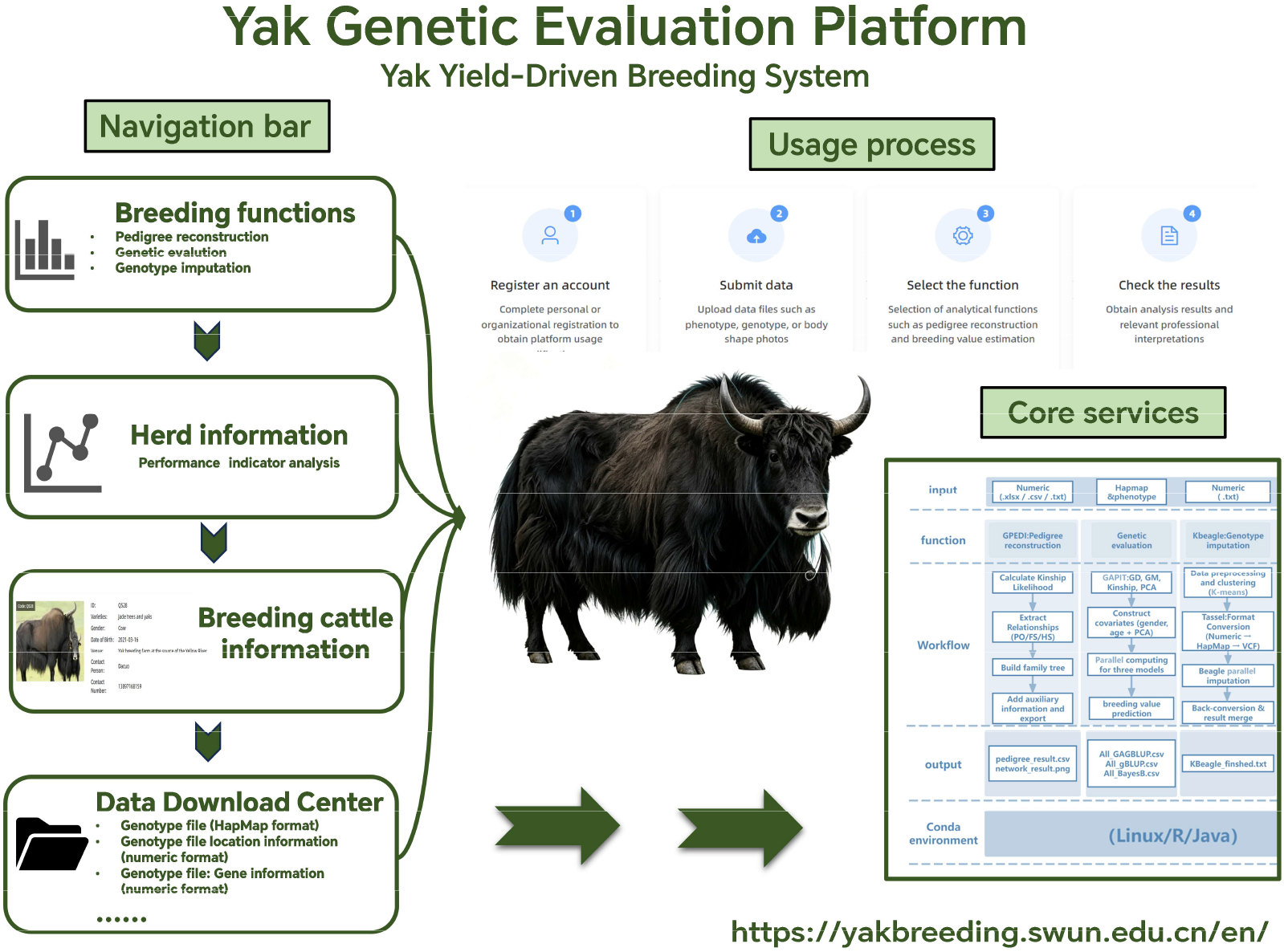
Schematic overview of the Yak Genomic Evaluation Platform (YGEP). Schematic overview of YGEP. The platform integrates user-end modules (data upload, analysis selection, result visualization) with a high-throughput bioinformatics backend, providing an integrated solution for yak genomic evaluation and breeding.

## 2. System design and implementation

YGEP was developed using a front-end and back-end separated Web architecture and integrated with automated analysis pipelines to provide yak genome evaluation and intelligent breeding functions. The platform was deployed on a CentOS Linux 8 server with an AMD EPYC 7763 processor (256 CPU cores). The front-end was developed based on the Vue.js framework and combined with the Element UI 2 component library to build the user interface. It runs in a Node.js 22 environment and is mainly responsible for user management, data uploading, task status monitoring, and visualization of analysis results **(Fig. S1)**. The back-end was developed using Java (JDK 11) and the Spring Boot 2 framework, which manages business logic, data processing, analysis task scheduling, and the execution of computational workflows. MySQL 8.0 was used for structured data storage, while Redis 6.x was integrated for cache management and task status tracking, improving system efficiency and stability.

YGEP adopts a user-specific data management strategy, in which each registered user is assigned an independent data storage space. Different functional modules are connected to separate data processing directories, and uploaded files are automatically stored in the corresponding module folders. To optimize bandwidth and storage efficiency, users are encouraged to upload datasets larger than 50 MB in compressed formats (e.g., .zip, .tar.gz). A background file monitoring service continuously scans user directories. When new input files are detected, the system automatically decompresses archives (if applicable), identifies the data type and initiates the corresponding analysis pipeline. After the analysis is completed, users can view or download results directly through the Web interface.

## 3. Utility and content

### 3.1 The yak genome database

Currently, the platform integrates phenotypic records from 855 individuals of 4 representative local yak populations **(Table S1)**. The data cover high-altitude yak core production areas such as Qinghai, Sichuan, and Xizang, including Yushu yak (298 individuals, accounting for 34.9%), Maiwa yak (281 individuals, accounting for 32.9%), Huanhu yak (183 individuals, accounting for 21.4%), and the Gaize yak (93 individuals, accounting for 10.9%). Among them, matched genotypic and phenotypic datasets from 706 individuals across three domestic populations (excluding the Gaize yak) were utilized for the experimental evaluations and module validations in this study. All corresponding phenotypic and genotypic datasets can be accessed and downloaded at https://yakbreeding.swun.edu.cn/en/download-file. Together, these datasets providing rich data support for cross-population genetic assessment and breeding analysis.

### 3.2 Core analytical modules

#### 3.2.1 Pedigree Reconstruction Model

To address the challenges of missing or inaccurate pedigree records in high-altitude grazing environments, YGEP integrates the GPEDI software (https://github.com/WangShikai101/GPEDI) for automated pedigree reconstruction and inbreeding analysis. The module accepts a numerical SNP genotype matrix encoded as 0/1/2 as input and automatically executes the GPEDI pipeline through the backend service. The pipeline infers pairwise genetic relationships from genome-wide SNP data and reconstructs pedigree information, including parent–offspring (PO), full-sib (FS), and half-sib (HS) relationships. After analysis, YGEP automatically generates standardized pedigree reconstruction files (pedigree_result.csv), individual inbreeding coefficient reports (inbreeding_result.csv), pedigree lineage tracing data (Tline_result.csv) and a network topology dataset (network_final.json). Subsequently, the backend processes this network JSON file using Cytoscape (v3.10.4) to automatically format and render a high-resolution, vector-based pedigree topology network (GPEDI_pedigree_result.svg) [14]. The backend also generates a vector-based pedigree topology network, which can be directly accessed and interactively explored through the Web interface, enabling intuitive visualization of population pedigree structure and genetic relationships **(Fig. 2A)**.

**Fig. 2.**
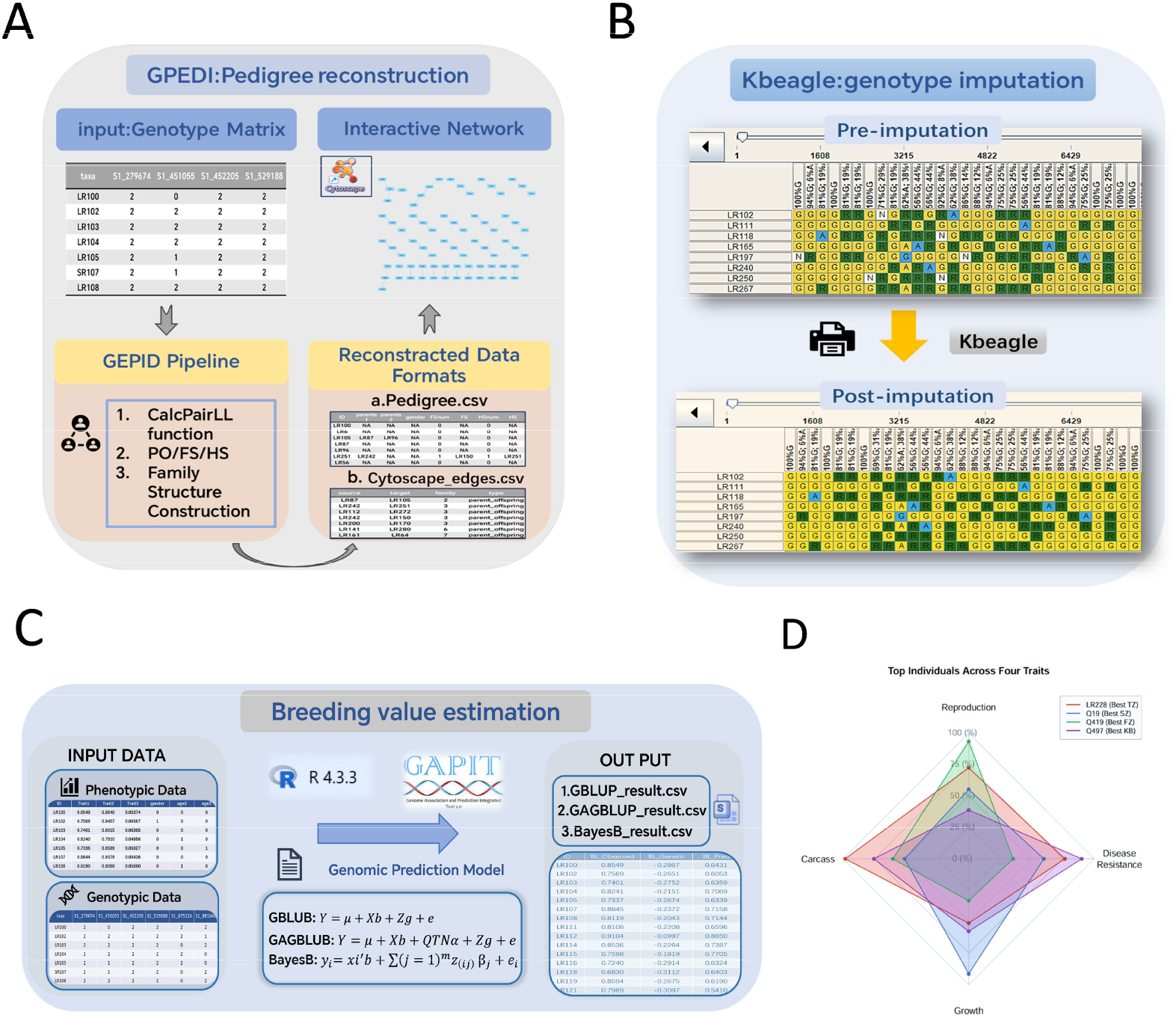
Functions of Yak Genetic Evaluation Platform. (A) Pedigree Reconstruction. (B) Genotype Imputation. (C) Breeding Value Estimation. (D) Radar chart of elite individuals evaluated for four key traits.

#### 3.2.2 Genotype Imputation

To reduce the economic cost of large-scale population genomic genotyping, YGEP integrates the KBeagle efficient parallel imputation algorithm [15]. This module automatically scans and identifies missing markers in low-density genotype datasets, including loci represented as NA or missing values. During computation, the KBeagle engine applies a pre-clustering strategy to optimize haplotype grouping structures, while multi-threaded parallel computation is implemented using the R package “parallel” to accurately infer and impute missing genotypes. The completed high-density genotype matrix (KBeagle_result.txt) is then automatically generated, providing high-quality genotype data with complete marker information for downstream analyses **(Fig. 2B)**.

#### 3.2.3 Breeding Value Estimation Model

The genetic evaluation module of YGEP was developed based on the R (v4.3.3) and the Genome Association and Prediction Integrated Tool (GAPIT v4) [16], leveraging multi-threaded parallel processing to accelerate automated GEBVs calculation. This module integrates individual phenotype and genotype data, fits fixed effects such as sex and age, and automatically extracts principal component analysis (PCA) eigenvectors as covariates to correct for population stratification.

For genomic prediction, the system incorporates three widely used and complementary prediction models to accommodate traits governed by different genetic architectures. The first is the Genomic Best Linear Unbiased Prediction (GBLUP) model [17], which assumes that SNP markers collectively contribute to genetic variation through genome-wide additive effects. The second is the Genomic Association-assisted Best Linear Unbiased Prediction (GAGBLUP) model [16], which integrates significant quantitative trait nucleotides (QTNs) identified by genome-wide association studies (GWAS) as fixed effects to improve prediction accuracy for traits influenced by major-effect loci. The third is the BayesB model, which is suitable for complex genetic architectures involving a mixture of small-effect polygenic variants and major-effect loci [18]. By combining these options, YGEP ensures optimal predictive performance across diverse breeding traits. After analysis, the platform automatically generates standardized output reports containing individual phenotypic information, genomic effects, and predicted breeding values, including GBLUP_result.csv, GAGBLUP_result.csv, and BayesB_result.csv **(Fig. 2C)**.

#### 3.2.4 Multi-trait breeding assessment and decision-making

The multi-trait comprehensive evaluation module was developed to eliminate scale differences among different traits. The system automatically reads breeding value files for each trait, performs Z-score normalization on the estimated genetic effects, and converts them into relative population percentiles (0–100%). Subsequently, the system calculates a comprehensive genetic index by averaging the percentile scores across multiple traits. Candidate superior breeding animals are ranked in descending order based on the index value. The Web interface simultaneously converts the multi-trait percentile data into radar charts **(Fig. 2D)**, providing an intuitive visualization of the comprehensive genetic potential distribution of candidate individuals and supporting breeding selection decisions.

## 4. Performance Evaluation

To evaluate the computational accuracy, scalability, and imputation performance of YGEP, systematic benchmark tests were conducted on the core computational modules integrated into the platform **(Fig. 3)**.

**Fig. 3.**
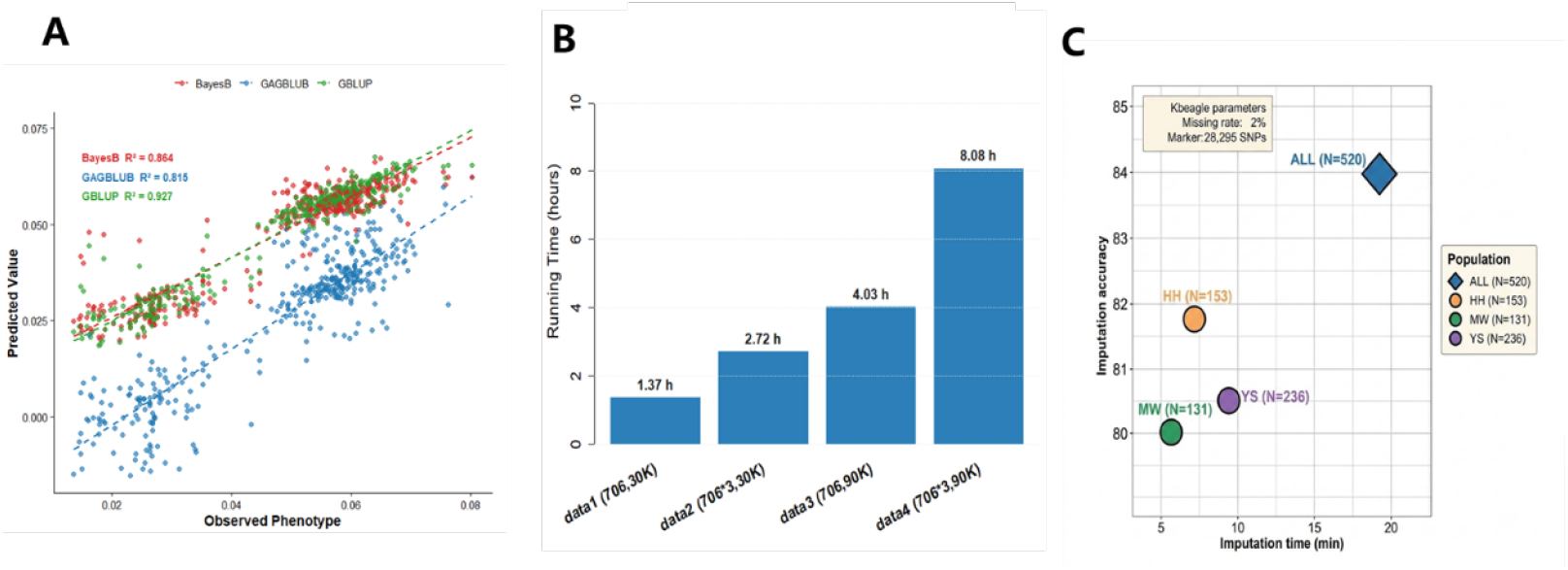
Performance evaluation of YGEP. (A) Genomic prediction performance of GBLUP, GAGBLUP, and BayesB for CCB in 706 yaks. (B) Computational runtime of YGEP under different sample sizes and marker densities. (C) Relationship between genotype imputation accuracy and runtime of KBeagle across three yak populations and their combined reference population.

Using a reference population consisting of 706 yaks, the prediction accuracy of GEBVs for the CCB trait was evaluated using three genomic prediction models (GBLUP, BayesB, and GAGBLUP). The correlation between GEBVs and observed phenotypes (r ± SE) were assessed **(Table S2)**. The results showed that GBLUP achieved the highest prediction accuracy, with a Pearson correlation coefficient of 0.96 ± 0.01 (R^2^ = 0.93). BayesB exhibited the second-best performance, with an accuracy of 0.93 ± 0.02 (R^2^ = 0.86), while GAGBLUP achieved an accuracy of 0.90 ± 0.02 (R^2^ = 0.82). These results demonstrate that GBLUP provides superior fitting stability and prediction performance for traits controlled by highly polygenic architectures in the yak population **(Fig. 3A)**.

Computational scalability tests confirmed robust linear performance across dataset sizes (**Fig. 3B)**. Execution times scaled predictably from 1.37 h for the baseline dataset (data1: 706 samples × 30K SNPs) to 2.72 h with a three-fold sample increase (data2: 2,118 × 30K), 4.03 h with a three-fold marker density increase (data3: 706 × 90K), and 8.08 h when scaling both dimensions simultaneously (data4: 2,118 × 90K), demonstrating YGEP’s capability to handle high-throughput genomic data efficiently.

Genotype imputation via the integrated KBeagle algorithm (28,295 SNPs, 2% missing rate) yielded accuracies of 80.0%-81.8% (runtime: 5.8-9.5 min) across individual populations (HH, MW, YS; N=131-236) **(Fig. 3C)**. Expanding the reference panel to the combined population (N=520) boosted imputation accuracy to 84.0% with a reasonable runtime of 19.3 min, validating the efficiency and cross-population applicability of KBeagle within YGEP.

## 5. Discussion

The main contribution of YGEP lies in unifying established genomic resources and tools into a continuous, application-oriented framework for yak genomic evaluation. While YGD provides genomic multi-omics data, and tools like GAPIT, GPEDI, and KBeagle handle specific analytical tasks, their practical application is often hindered by fragmented software environments and tedious data format conversions [19]. YGEP bridges these gaps by streaming pedigree reconstruction, genotype imputation, genomic prediction, and multi-trait evaluation into an automated workflow, eliminating manual intermediate file transfers. Beyond workflow integration, YGEP substantially improves overall computational throughput. Instead of sequentially executing GAPIT for each phenotype, YGEP employs R-based multi-threaded parallel processing to evaluate multiple traits concurrently. Similarly, for genotype imputation, KBeagle partitions genotypic data into subpopulations via K-means clustering and executes multi-threaded imputation across these clusters in parallel, significantly accelerating processing efficiency in genetically differentiated yak populations. Together, these high-throughput optimizations allow YGEP to serve as a practical, scalable decision-support system that lowers technical barriers for implementing genomic selection in yak breeding.

The genomic prediction results further demonstrate the applicability of YGEP for genetic evaluation in yaks. GBLUP achieved the highest predictive accuracy for CCB (r = 0.96), followed by BayesB (r = 0.93) and GAGBLUP (r = 0.90). The high performance of GBLUP is consistent with a predominantly polygenic genetic architecture of CCB, in which many loci with small effects contribute to genetic variation. In Hanwoo cattle, GBLUP has also shown strong predictive performance for growth and carcass traits, particularly when genetic variation is distributed across multiple loci rather than dominated by major QTL. Similarly, comparable performance of GBLUP and BayesB has been reported for growth and carcass traits in Brangus cattle [20]. The higher accuracy of GBLUP in the present study therefore suggests that genome-wide additive relationships captured more genetic signal than heterogeneous marker effects. The lower accuracy of GAGBLUP may reflect the limited contribution of the selected significant QTNs to CCB genetic variance, consistent with recent findings that incorporating GWAS information does not necessarily improve genomic prediction when major-effect loci are limited [21]. GAGBLUP treats GWAS-identified QTNs as fixed effects; when the trait is highly polygenic and no single QTN explains a large proportion of variance, this specification may introduce noise rather than improve prediction.

Genotype imputation is an important component of YGEP because it can increase marker density while reducing genotyping costs. In the present study, KBeagle achieved 80.0–81.8% accuracy within individual populations and 84.0% using the combined reference panel, indicating that reference-panel size and genetic diversity affect imputation performance. Previous studies using Beagle(v5.4) have reported higher accuracies under favorable reference-panel and marker-density conditions [22], while KBeagle has shown improved accuracy and computational efficiency in genetically differentiated yak populations. While the 84.0% accuracy is promising for routine genomic evaluations, applications requiring highly precise individual genotypes may necessitate further optimization. Further evaluation of its effects on genomic relationships and GEBV accuracy will be important for practical breeding applications.

Despite these advantages, several limitations of YGEP remain. The current version supports only four predefined yak populations, which may not adequately represent the genetic diversity of yaks across regions and production systems. Expanding the reference populations will improve the robustness of genotype imputation and genomic evaluation. In addition, phenotypic data are provided by users since downstream analyses heavily depend on the completeness and quality of the uploaded records. Standardized phenotype formats and quality-control procedures will therefore be needed to improve data reliability. Finally, YGEP currently focuses on genotype-, phenotype-, and pedigree-based evaluation and does not incorporate multi-omics data. Integrating transcriptomic, proteomic, and epigenomic information in future versions could provide additional biological evidence for interpreting candidate loci and economically important traits.

Future development of YGEP will focus on expanding its data coverage and analytical capabilities. Increasing the size and diversity of the reference population will improve the transferability of genotype imputation and genomic prediction across yak populations. To address the reliance on user-provided data quality, future updates will introduce an automated quality control (QC) module to detect phenotypic anomalies, filter outliers, and standardize formats prior to downstream analyses. More genomic prediction models and trait-specific strategies could also be incorporated to accommodate different genetic architectures and breeding objectives. Integrating multi-omics resources would further facilitate the functional interpretation of genomic signals and candidate genes. In addition, artificial intelligence and machine-learning methods could be used to integrate these data and assist animal ranking, selection, and breeding decisions. These improvements would further enhance YGEP as a practical decision-support platform for precision yak breeding.

## 6. Conclusion

This study developed the YGEP, aiming to promote digital herd management, high-throughput phenotypic measurement, and genomic selection in yak breeding. YGEP provides an automated analysis pipeline including pedigree reconstruction, genotype imputation and multi-model GEBV estimation, achieving a closed-loop processing from raw field data to precise breeding decisions. In the future, the platform functions, analysis algorithms, and data capacity of YGEP will continue to be updated and expanded, incorporating larger reference populations to assist breeding personnel and researchers in efficiently implementing precision breeding in grazing environments. YGEP will provide a valuable technical platform for researchers and breeders engaged in yak genetic evaluation, molecular breeding, and germplasm resource conservation.

## Abbreviations

YGEP: Yak Genomic Evaluation Platform
GEBVs: Genomic Estimated Breeding Values
GS: Genomic Selection
YGD: the Yak Genome Database
CCB: Cannon Bone Circumference
GAPIT: Genome Association and Prediction Integrated Tool
GBLUP: Genomic Best Linear Unbiased Prediction
GAGBLUP: Genomic Association-assisted Best Linear Unbiased Prediction
PCA: Principal Component Analysis
GWAS: Genome-Wide Association Study
QTN: Quantitative Trait Nucleotide

## Author contributions

**Y.Y**. contributed to Conceptualization, Methodology, Formal analysis, Investigation, Resources, Writing – original draft, Writing – review & editing, Visualization, and Supervision. **Z.L**. and X.S. contributed to Validation and Investigation. **Y.F**. contributed to Validation, Investigation, and Resources. **Z.Z**. contributed to Software, Data curation, and Validation. **X.G**. contributed to Software, Validation, and Formal analysis. **S.W**. contributed to Resources and Data curation. **J.Z. (Jiahong Zhao)** contributed to Formal analysis and Investigation. **J.Z. (Jincheng Zhong)** contributed to Software and Investigation. **L.S** and **D.L**. contributed to Conceptualization, Formal analysis, Investigation, Writing – original draft, Supervision, and Funding acquisition. **J.W**. contributed to Conceptualization, Methodology, Writing – review & editing, Visualization, Supervision, and Funding acquisition. All authors have read and agreed to the published version of the manuscript.

## Data availability

All related codes are accessible at the following URLs: https://github.com/yy0587023-rgb/platform. The Platform can be addressed at: https://yakbreeding.swun.edu.cn/en/. The platform will be maintained and updated for at least five years, with new reference populations and prediction models added periodically.

## Consent for publication

Not applicable.

